# Neurophysiological coding of statistical and deterministic rule information

**DOI:** 10.1101/2020.10.14.338913

**Authors:** Ádám Takács, Andrea Kóbor, Zsófia Kardos, Karolina Janacsek, Kata Horváth, Christian Beste, Dezső Németh

## Abstract

Humans are capable of acquiring multiple types of information presented in the same visual information stream. It has been suggested that at least two parallel learning processes are important during learning of sequential patterns – statistical learning and rule-based learning. Yet, the neurophysiological underpinnings of these parallel learning mechanisms in visual sequences are not fully understood. To differentiate between the simultaneous mechanisms at the single trial level, we apply a temporal EEG signal decomposition approach together with sLORETA source localization method to delineate whether distinct statistical and rule-based learning codes can be distinguished in EEG data and can be related to distinct functional neuroanatomical structures. We demonstrate that concomitant but distinct aspects of information coded in the N2 time window play a role in these mechanisms: mismatch detection and response control underlie statistical learning and rule-based learning, respectively, albeit with different levels of time-sensitivity. Moreover, the effects of the two learning mechanisms in the different temporally decomposed clusters of neural activity also differed from each other in neural sources. Importantly, the right inferior frontal cortex (BA44) was specifically implicated in statistical learning, confirming its role in the acquisition of transitional probabilities. In contrast, rule-based learning was associated with the prefrontal gyrus (BA6). The results show how simultaneous learning mechanisms operate at the neurophysiological level and are orchestrated by distinct prefrontal cortical areas. The current findings deepen our understanding on the mechanisms how humans are capable of learning multiple types of information from the same stimulus stream in a parallel fashion.

## 1. Introduction

How the human brain encodes regularities of the environment is a current topic in cognitive neuroscientific research. It has been proposed that learning of patterns includes at least two parallel processes (Batterink et al., 2015; Conway, 2020; Maheu et al., 2020; Nemeth et al., 2013a). One of them is called statistical learning and refers to an automatic acquisition of associative relations and frequencies of external stimuli (Conway, 2020). Another attention-dependent mechanism plays a role in controlling the learning process, learning of non-adjacent relations, and integrating rule-based regulations. This second process is often labelled as higher-order sequence learning (Howard and Howard, 1997; Nemeth et al., 2013a), deterministic rule-learning (Maheu et al., 2020), or order-based learning (Simor et al., 2019). It has been proposed that humans organise the temporal regularities in their environment in two distinct hypothesis spaces (Conway, 2020; Maheu et al., 2020): one is based on the estimation of probabilities, and works as statistical bias; the other one is based on deterministic relations or rules. Importantly, in situations where both statistical and rule-like regularities occur in parallel (e.g., in language or music), not only competition can occur but rules can induce statistical biases and vice versa (Maheu et al., 2020).

Parallel learning of different types of regularities can be studied with variations of sequence learning tasks, in which rules and inter-stimulus dependencies are repeated in the stimulus stream (Conway, 2020; Howard and Howard, 1997; Nissen and Bullemer, 1987). A short sequence can be memorised as an arbitrary order of items and stored in declarative memory (Conway, 2020). However, when the information stream becomes longer and more complicated, humans can only encode frequently co-occurring patterns (statistical information) and hierarchical structures within the sequences (rule-based learning) (Conway, 2020; Nemeth et al., 2013a). While statistical learning occurs in a rapid manner and reaches its plateau quickly, learning of sequential rules is characterized by a gradual change (Kóbor et al., 2018; Simor et al., 2019). Despite the growing interest in these simultaneous learning mechanisms (Conway, 2020), the neurophysiological processes behind them remain largely unknown (Kóbor et al., 2019, 2018) and it is an open question how these parallel learning mechanisms are coded at the neurophysiological level.

However, solving this puzzle represents a methodological challenge. Namely, different aspects of task-related information and spontaneous background activities are present simultaneously in the EEG recordings of the scalp (Folstein and Van Petten, 2008a; Nunez et al., 1991; Ouyang and Zhou, 2020; Stock et al., 2017) and overlapping brain areas are involved in processing them (Mückschel et al., 2017a). Furthermore, coding levels are more likely to get intermixed if the task requires manipulations of stimulus- and response-related representations at the same time, such as in interference suppression (Mückschel et al., 2017a) or stimulus-response binding (Opitz et al., 2020; Takacs et al., 2020b, 2020a). Using averaged trials of EEG data (Kóbor et al., 2018), it has been demonstrated that the frontal N2 event-related potential (ERP) component reflects a rapid automatic detection of statistical properties in the sequence, *and* a rule-based (pattern vs random) learning with a longer time course of development. The P3 ERP-component seems to be sensitive only to rule-based differences and not to statistical properties (Kóbor et al., 2018). In the EEG, the N2 is not a unitary component: it can be divided into different subcomponents related to cognitive control and stimulus-related detection of novelty information processing (Adelhöfer et al., 2018, 2019; Chmielewski et al., 2018; Folstein and Van Petten, 2008b; Mückschel et al., 2017a). Specifically, subprocesses in the N2 time window that are affected by stimulus-related information can be expected to show sensitivity to statistical learning. On the other hand, subprocesses related to control and monitoring may show modulations in relation to rule-based learning of sequential information. Since statistical learning and deterministic rule-based learning are supposed to occur in parallel (Conway, 2020), it is likely that these different information encoding principles are concomitantly coded in the neurophysiological signal. To solve this problem, we employed a signal decomposition method to disentangle the proposed simultaneous mechanisms.

A powerful method to disentangle intermixed coding levels in EEG data is the residue iteration decomposition (RIDE) (Chmielewski et al., 2018; Mückschel et al., 2017a, 2017b; Ouyang et al., 2015; Ouyang and Zhou, 2020). Instead of a traditional averaging of single trial data, RIDE can dissociate between three main activity clusters which preserve the dynamical response pattern of the single trial data. These activity clusters are the stimulus-related S-cluster, the response-related R-cluster, and finally the C-cluster which captures the not strictly perceptual or motor aspects of the signal. That is, the C-cluster reflects the translational aspect that is related to the association of stimuli with the appropriate response (Ouyang et al., 2017). The method postulates that parallel processes have different latency characteristics: perceptual and posterior attention processes are more likely tied to the stimulus presentation (S-cluster), while motor execution is locked to the response (R-cluster) (Ouyang et al., 2011). Furthermore, higher-order mechanisms, such as visuo-motor integration, memory, retrieval, and decision making have variable latencies (C-cluster), and, therefore, the related neurophysiological signals are smeared in the undecomposed EEG recording. In a previous study, different N2 effects related to conflict detection and response control could be differentiated by temporal signal decomposition (Mückschel et al., 2017a). Based on the parallel subprocesses notion of Conway (2020, see also Kóbor et al., 2018), it is expected that statistical learning as an automatic, stimulus-driven process is reflected by the S-cluster activity, while the attention-related, higher-order rule-based learning is mainly reflected by the C-cluster activity. If this is the case, the current study would be the first to relate cognitive distinctions of statistical and rule-based learning to distinct coding processes at the neurophysiological level.

Learning of sequential patterns has been tied to widespread activations including the parietal cortex, the prefrontal cortex (PFC), the hippocampus, the cerebellum, and the basal ganglia (Conway, 2020). Yet, at least two networks can be differentiated from each other: a more perceptual, posterior one, and a prefrontal one (Conway, 2020). Especially the lateral PFC is important in the processing of rule-based information in temporal sequences (Conway, 2020; Janacsek et al., 2015). Prefrontal functions, such as response selection and selective attention likely play a role not only in the production of sequential action but also in learning of sequenced information (Conway, 2020). For instance, learning of non-adjacent, long-distance regularities involves the inferior frontal gyrus (IFG) (Barascud et al., 2016; Conway, 2020; Maheu et al., 2020; Southwell and Chait, 2018). Specifically, the left IFG has been proposed as a supra-modal hierarchical processor of sequence information (Tettamanti and Weniger, 2006), especially when sequences consisted of grammar-like structures or verbal stimuli (Maheu et al., 2020). However, processing sequences of visuospatial or visuomotor items seem to show right hemisphere dominance (Janacsek et al., 2015; Jarret et al., 2019; Roser et al., 2011) and data from brain stimulation experiments have shown that the right dorsolateral prefrontal cortex (DLPFC), but not the left DLPFC is associated with learning and retention of visuomotor sequences (Janacsek et al., 2015). However, prefrontal functions, such as attention and inhibitory control have been suggested to have an orthogonal relationship with statistical learning (Conway, 2020; Filoteo et al., 2010; Nemeth et al., 2013b). That is, performance on prefrontal functions negatively correlates with statistical learning abilities; moreover, temporary decrease of executive functions can enhance statistical learning (Ambrus et al., 2020; Nemeth et al., 2013b; Virag et al., 2015).

The controversy between the role of frontal neural activity, particularly in the IFG in statistical learning and the opposing relation between prefrontal executive functions and learning of probabilistic information can be solved by considering alternative functions for the right IFG (Erika-Florence et al., 2014). Specifically, the alternative attentional rule-processing (ARP) account of the right IFG postulates that this structure is not specific for inhibitory (or other executive) functions, but houses general task-related functions, such as detecting novelty and frequency information in task settings, and gating of learning through attention control (Erika-Florence et al., 2014; Southwell and Chait, 2018). Thus, the orthogonal relationship between statistical learning and prefrontal executive functions does not exclude the possibility of the involvement of IFG in learning, especially when the sequence is presented visually. In such sequences, detection of mismatch between internal models of the sequence and surprising new items in the information stream contributes to the calculation of uncertainty. This uncertainty or “surprise” detection has been linked to the right IFG (Barascud et al., 2016; Southwell and Chait, 2018). In the current study, statistical learning effects are studied by temporally decomposed ERP components. Among these components, the S-cluster N2 is expected to show the mismatch function related to distinguishing between predictable (frequent) and unpredictable (rare) stimuli. Since learning of statistical regularities is functionally related to detecting novelty in the stimuli, based on the ARP model, we expected that learning of statistical probabilities as an S-cluster N2 effect is related to the right IFG (Barascud et al., 2016; Conway, 2020; Maheu et al., 2020; Southwell and Chait, 2018). In contrast, we expected more widespread prefrontal activations related to rule-based learning effect in the C-cluster N2 in accordance with previous brain stimulation studies (Conway, 2020). In sum, to dissociate between neurophysiological markers of parallel learning mechanisms within sequence learning, we employed temporal signal decomposition and subsequent source localization of the RIDE-decomposed components.

## 2. Methods

### 2.1. Participants

The current study is a re-analysis of the sample of 40 undergraduate students (21.4 years ± 1.6) as reported in Kóbor et al. (2018). All of the participants reported their vision to be normal or corrected-to-normal, the lack of any neurological or psychiatric condition, or taking any psychoactive medication. Further details of the participants, including handedness, years of education, and performance on standard neuropsychological tests are reported in the original study (Kóbor et al., 2018).

### 2.2 Ethics statement

Before the experiment started, participants were informed about the procedures of measurement and data collection. All participants provided written informed consent prior to participating, for which they received either payment or course credit. The study received approval from the United Ethical Review Committee for Research in Psychology in Hungary (EPKEB). The experiment was conducted in accordance with the Declaration of Helsinki.

### 2.3 Task

To measure sequence learning, a modified version of the Alternating Serial Reaction Time (ASRT) task (Howard and Howard, 1997) was presented to the participants, while EEG was recorded. This variation of the task is called the cued ASRT (Nemeth et al., 2013a) and it has been successfully adapted for EEG measurement before (Kóbor et al., 2018). In this version of the task, a target stimulus (either a black or a red arrow facing left, up, down, or right) was presented on the display. The stimuli were always arranged to a central position of the screen. The task’s instruction asked for pressing the corresponding button on the response box (indicating the four possible directions, see Figure 1) as accurately and as fast as possible. The stimuli appeared according to an 8-element alternating sequence. The alternating sequence determined that random elements were always followed by pattern elements, and vice versa. Following this rule of stimulus presentation, if the sequence was 1-2-4-3, the stimuli appeared as 1-r-2-r-4-r-3-r, where the four numbers correspond to the arrow’s possible directions and ‘r’ stands for a random direction. To visually indicate the task’s rule, the black arrows represented pattern elements and the red arrows represented random elements. The instruction explicitly stated that the black arrows always followed a pattern, while red arrows appeared in random directions. Thus, the sequential rule was made explicit beforehand for the participants, and this information stayed salient during the task with the use of different colours for the two types of stimuli. Participants were asked to find the pattern in how the black arrows were presented to perform better in the task.

**Figure 1.**
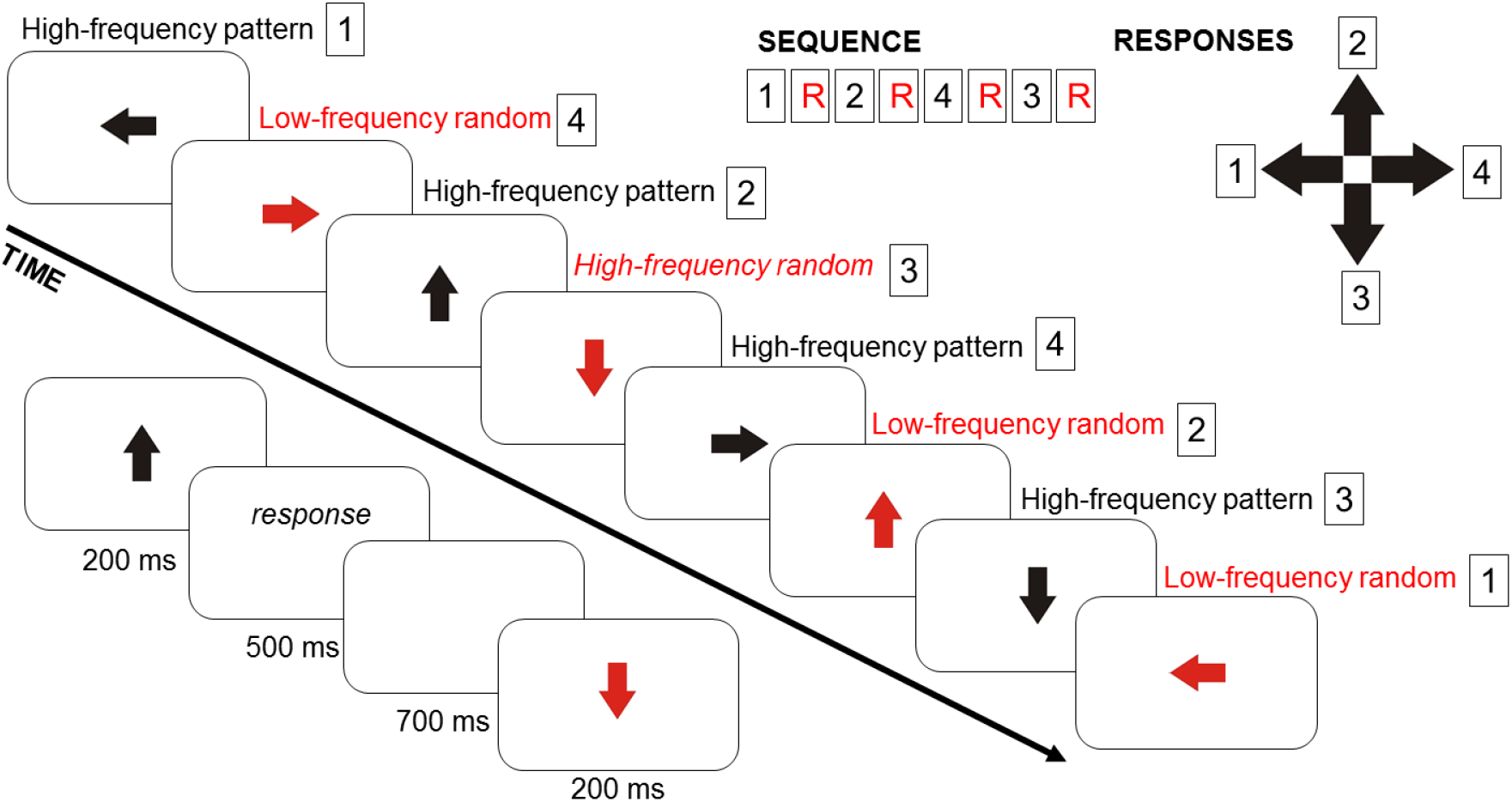
Experimental design. In the current version of the Alternation Serial Reaction Time (ASRT) task, participants saw an arrow in the middle of the screen. The arrows’ presentation followed an eight-element sequence, in which pattern and random (R) elements alternate. Regularity of the sequence (e.g., the task rule) has been marked by black colour for pattern and red colour for random elements. Numbers denote the four possible directions of the arrows (1 – left, 2 – up, 3 – down, 4 – right). These directions correspond to the configuration of the response pad as presented in the top right corner of the figure. Some series of consecutive elements (triplets) occur frequently in the task than others. High-frequency triplets could either end with pattern or with random elements, while low-frequency triplets always end with a random element. Timing of the task is presented in the left bottom corner of the figure.

The structure of alternating black and red arrows means that some three-element series of consecutive stimuli (‘triplets’) appeared with different levels of probabilities, that is, triplets with high and low frequencies could be identified. In the example of 1-r-2-r-4-r-3-r, the combinations of 1-X-2, 2-X-4, 4-X-3, 3-X-1 (X stands for any middle stimulus within the triplet) are frequent ones. However, series as 3-X-2 or 4-X-2 are less common. In these latter examples, the third element of the triplet can never occur as part of the sequence (pattern element), while in the former, frequent ones, the triplet can end either as a pattern or as a random stimulus. Triplets that are more probable to occur are called high-frequency triplets, while less probable ones are called low-frequency triplets. The designations also refer to the transition probabilities within the triplets. That is, in a high-frequency triplet, the third element is highly predictable based on the first element (with 62.5% probability). In the case of the low-frequency triplet, the predictability of the last element is lower (12.5%). In addition, each element can be categorized according to its sequential position, i.e., whether it is a pattern element or a random element. If the last element is a pattern one, the explicit knowledge of the sequential rule contributes to predicting the direction of the next arrow. Therefore, the combination of the alternating sequence structure and the frequency information of the triplets results in a threeway categorization of regularities within the task: high-frequency triplets with either pattern or random endings and low-frequency triplets that always end with random elements. Consequently, based on the frequency and structure of the triplets, three types of trials can be distinguished: (1) high-frequency pattern trials; (2) high-frequency random trials, and; (3) low-frequency random trials. These types of trials can be analysed to track the course of the two most important learning mechanisms in the ASRT task: sequence or rule-based learning and statistical learning. Learning of sequential rule is quantified as a difference in response times between high-frequency pattern and high-frequency random elements. These elements are the last parts of a high-frequency triplet, thus, they represent the same frequency information across the task. However, they differ in terms of sequential position, as one is part of the reoccurring pattern, while the other appears randomly. Therefore, faster responses to the pattern compared to the random trials indicates better learning of the sequential rule. To calculate the acquisition of frequency-based information, that is, statistical learning, response times for high-frequency random and low-frequency random trials are compared. Here, the elements carry the same information of the sequential rule (both random) but differ in frequencies, since they appear either in a high-frequency or a low-frequency triplet. Therefore, faster response time to high-frequency random elements than to low-frequency random elements is considered as a behavioural marker for successful statistical learning. In summary, statistical learning captures purely probability-based learning, while learning of the sequential rule captures order-based learning. For the detailed structure and timing of the experiment, see Figure 1 and as reported in Kóbor et al. (2018).

### 2.4 EEG recording and analysis

Scalp EEG was recorded with 64 Ag/AgCl electrodes from an elastic cap (EasyCap, Germany), using Synamps amplifier and Neuroscan recording software (Compumedics Neuroscan, USA). For reference and ground, the tip of the nose and AFz electrode were employed. EEG recording was performed with a sampling rate of 1000 Hz and with an online filter of 70 Hz low-pass, 24 dB/oct. Impedance levels of electrodes were kept below the threshold of 10 kΩ. The recordings were analysed in BrainVision Analyzer 2.0 (Brain Products, Germany). The EEG data were band-passed filtered (0.5-30 Hz, 48 dB/oct), and a notch filter was applied (50 Hz). Then, independent component analysis was performed to remove components which were identified as eye-movement artifacts and heart beats based on their temporal properties and spatial distributions (Delorme et al., 2007). Then, EEG data were re-referenced to the average of all electrodes. The pre-processed EEG was segmented according to time-on-task and experimental conditions, which required two consecutive steps. First, six equal length time bins or epochs were created. Second, the data were segmented into the three experimental conditions in each epoch. That is, high-frequency pattern, high-frequency random, and low-frequency random segments were created. Segments with incorrect responses or without response markers (misses) were not included in the segmentation to ensure that only those trials are analysed which were correctly identified by the participants. These segments were 800 ms long, starting −200 ms before the stimulus presentation and ending 600 ms after that. Following the segmentation, automatic artefact rejection (as implemented in BrainVision Analyzer 2.0) was used to remove the remaining artifacts (with a voltage threshold of +/− 100 μV at any channels). The kept segments were then baseline corrected based on the activity in a 200-ms-long interval before the stimulus presentation. The final segments represented the experimental conditions in six consecutive time stages of the task with correct responses. Further details of the EEG processing, including justification of these steps, can be found in Kóbor et al. (2018). The original analysis identified a frontal N2 component (time window of 200-300 ms) and a P3 component with a maximum activity on the electrode Pz (time window of 250-350 ms). Results of the original ERP analysis are reported in Kóbor et al. (2018). In the current study, the preprocessed, segmented, cleaned, and baseline-corrected data was re-analyzed with RIDE temporal decomposition.

### 2.5 Residue iteration decomposition

The RIDE temporal decomposition method was used for trial-to-trial variability-based analysis of the stimulus-locked ERPs. RIDE aims to distinguish components with variable intercomponent delays based on the single trial latency variability information (Ouyang et al., 2015; Ouyang and Zhou, 2020). The single-trial ERPs are decomposed into different components with differential latency variability. Temporal decomposition is achieved in an iterative way. RIDE assesses latency variability in a channel-specific fashion, thus, differences between individual electrodes remain valid in the identified components (Ouyang et al., 2015). In the current study, we used the RIDE toolbox in Matlab (Mathworks, Inc., Massachusetts, USA), and followed the protocols of earlier studies (Mückschel et al., 2017a; Ouyang et al., 2011; Verleger et al., 2014). For a review of previous RIDE applications, please see Ouyang and Zhou (2020). In two cluster types, latency information was extracted based on existing marker information. That is, stimulus onset was used to create the S-cluster (“stimulus cluster”) and response markers for the R-cluster (“response cluster”). The C-cluster (“central cluster”) was estimated and iterated in each trial, assuming a non-constant latency. That is, virtual time markers were created to decompose the last cluster. RIDE uses an iterative decomposition with an *L1*-norm minimization. The resulting median waveforms represent the dynamics of single trials more reliably than traditional averaging of ERP components (Ouyang et al., 2017; Ouyang and Zhou, 2020). To estimate the first decomposed cluster, RIDE subtracts the other two from each trial and adjusts the residual of all trials for the latency information of the first one. As a result, median waveform is created for all time points in the cluster’s search interval. The procedure is then repeated to create the remaining two clusters. The whole process iterates until convergence to better estimate the sub-components. To extract the waveforms of each cluster, search windows should be predefined (Ouyang et al., 2015, 2011). The following time-intervals were specified: for the S-cluster up to 500 ms after the stimulus presentation; for the R-cluster 300 ms before and after the correct response marker; for the C-cluster 150 ms to 600 ms after the stimulus. Based on the original study (Kóbor et al., 2018), we selected the electrode Fz for the N2 component, and the electrode Pz for the P3 component. The selected channels were then visually inspected to determine the time window of these two components separately for the three RIDE clusters. The N2 component was identified between 200 and 300 ms after the stimulus presentation in the S-cluster (similar to the undecomposed N2 in Kóbor et al. (2018). In the C-cluster, the N2 was visible between 240 and 340 ms after the stimulus onset. In the R-cluster, the N2 could not be identified by visual inspection, however, it was analysed in the time window corresponding to the undecomposed N2 and the S-cluster N2 time window (200-300 ms). The original study reported the P3 component between 250 and 350 ms after the stimulus presentation (Kóbor et al., 2018). We selected the identical time window in the S-cluster data. In the C-cluster, the P3 was detected between 250 and 400 ms, and in the R-cluster, between 280 and 440 ms after the stimulus onset. Within the selected time intervals, the mean amplitude was quantified and extracted at the single-subject level.

### 2.6 Source localization

The standard low resolution brain electromagnetic tomography (sLORETA) (Pascual-Marqui, 2002) was used to examine the estimated neural sources of the effects of statistical learning and rule learning for the temporally decomposed EEG data. sLORETA provides neural source information based on images of standardized current source density. The standard electrode coordinates according to the 10/20 system were used as input. Then, a three-shell spherical head model and the covariance matrix were calculated using the baselines at the single subject level. Within the head model, the intra-cerebral volume is partitioned into 6239 voxels using a spatial resolution of 5 mm. The standardized current density is calculated for each voxel, using an MNI152 head model template. sLORETA provides a single linear solution for the inverse problem without localization bias (Marco-Pallarés et al., 2005; Pascual-Marqui, 2002; Sekihara et al., 2005). Sources identified with using the sLORETA have been validated in combined MRI/EEG and TMS/EEG studies (Dippel and Beste, 2015; Ocklenburg et al., 2018; Sekihara et al., 2005). Comparisons against zero were used for the sLORETA contrasts. To calculate the statistics on the sLORETA sources, we used voxel-wise randomization tests with 2500 permutations and statistical nonparametric mapping procedures (SnPM). Locations of voxels that were significantly different *(p* < .05) are shown in the MNI-brain. Significant activations represent critical t-values corrected for multiple comparisons as implemented in the sLORETA software.

### 2.7 Statistics

Statistical analyses were performed by using IBM SPSS Statistics, and followed the established procedure of analyzing the ASRT task (Kóbor et al., 2018; Nemeth et al., 2013a; Song et al., 2007). The two main learning processes, statistical learning and rule-based learning were quantified for the behavioural and EEG analyses. Statistical learning was defined as the difference between high-frequency random and low-frequency random trials in reaction time, and mean activity of the RIDE clusters. Better statistical learning means shorter reaction time for high-frequency random than for low-frequency random trials. This learning index is expected to become larger as the learning progresses. Rule-based learning was quantified as a difference between high-frequency pattern and high-frequency random trials in reaction time, and mean activity of the RIDE clusters. Better rule-based learning means shorter reaction time for the pattern than for high-frequency random trials. This learning index also becomes larger as the learning progresses (revealed by a significant interaction with the epoch). The two learning mechanisms were analyzed in two-way repeated measures ANOVAs with “type” (high-frequency pattern, high-frequency random, and low-frequency random) and “epoch” (from one to six) as within-subject factors on reaction time, and the mean amplitude of the N2 and P3 components in the three RIDE clusters. When the interaction between type and epoch was significant, statistical learning was tested with a type by epoch ANOVA, in which the type factor included high-frequency random and low-frequency random trials. Similarly, rule-based learning was analyzed as a type by epoch ANOVA, in which the type factor included high-frequency random and high-frequency pattern trials. In these ANOVA models, the Greenhouse-Geisser epsilon correction was used when the lack of sphericity necessitated it. Effect sizes are reported as partial eta-squared. Post-hoc pairwise comparisons of the decomposed N2 and P3 mean amplitudes were Bonferroni-corrected, if necessary.

## 3. Results

### 3.1 Behavioural results

Details of the behavioural results, including main effects and descriptive data are reported and illustrated in Kóbor et al. (2018). Here we summarize only those behavioural results that indicate learning effects and are necessary to interpret the neurophysiological results. Importantly, the type (high-frequency pattern, high-frequency random, and low-frequency random) by epoch (1-6) ANOVA on the reaction time data showed a significant type by epoch interaction (*F*(10, 390) = 15.25, ε = .382,*p* < .001, ηp^2^ = .281). This indicates that participants’ responses changed between triplet types during the task. In the case of rule-based learning, the type by epoch ANOVA revealed a significant type by epoch interaction (*F*(5, 195) = 16.78, ε = .617, *p* < .001, ηp^2^ = .301). Participants’ responses were faster in high-frequency pattern than in high-frequency random condition in all six epochs *(p* < .001), and this difference gradually increased from the first to the fourth epoch (smaller in the 1^st^ epoch than in the other epochs *p*s < .001; smaller in the 2^nd^ epoch than in the 4^th^, 5^th^, and 6^th^ ones *p* < .008; smaller in the 3^rd^ epoch than in the 4^th^, 5^th^, and 6^th^ ones*p* < .021; and smaller in the 4^th^ epoch than in the 6^th^*p* = .035; for descriptive data, see Kóbor et al. (2018). That is, participants showed rule-based learning from the beginning of the task, and this learning effect became larger as the task progressed. In the case of statistical learning, the type by epoch ANOVA did not reveal a significant type by epoch interaction *(p* = .643), however, the main effect of type was significant (*F*(1, 39) = 123.53,*p* < .001, ηp^2^ = .760). Participants were faster in high-frequency random than in low-frequency random condition, throughout the task. That is, while rule-based learning showed a gradual adaptation to the task, sensitivity to statistical regularities developed quickly and remained stable.

### 3.2 Neurophysiological results

Analyses of the decomposed N2 and P3 components are presented as follows. First, type by epoch repeated measure ANOVAs are described. Importantly, post-hoc analyses were only conducted in case of significant type or type by epoch effects. These post-hoc analyses are separate ANOVAs investigating statistical learning or rule-based learning.

#### 3.2.1 Decomposed N2 (S-cluster)

Grand-averages of ERP waveforms in the S-cluster N2 time window split by triplet type and epoch are presented in Figure 2 and statistical results are summarized in Table 1.

**Figure 2.**
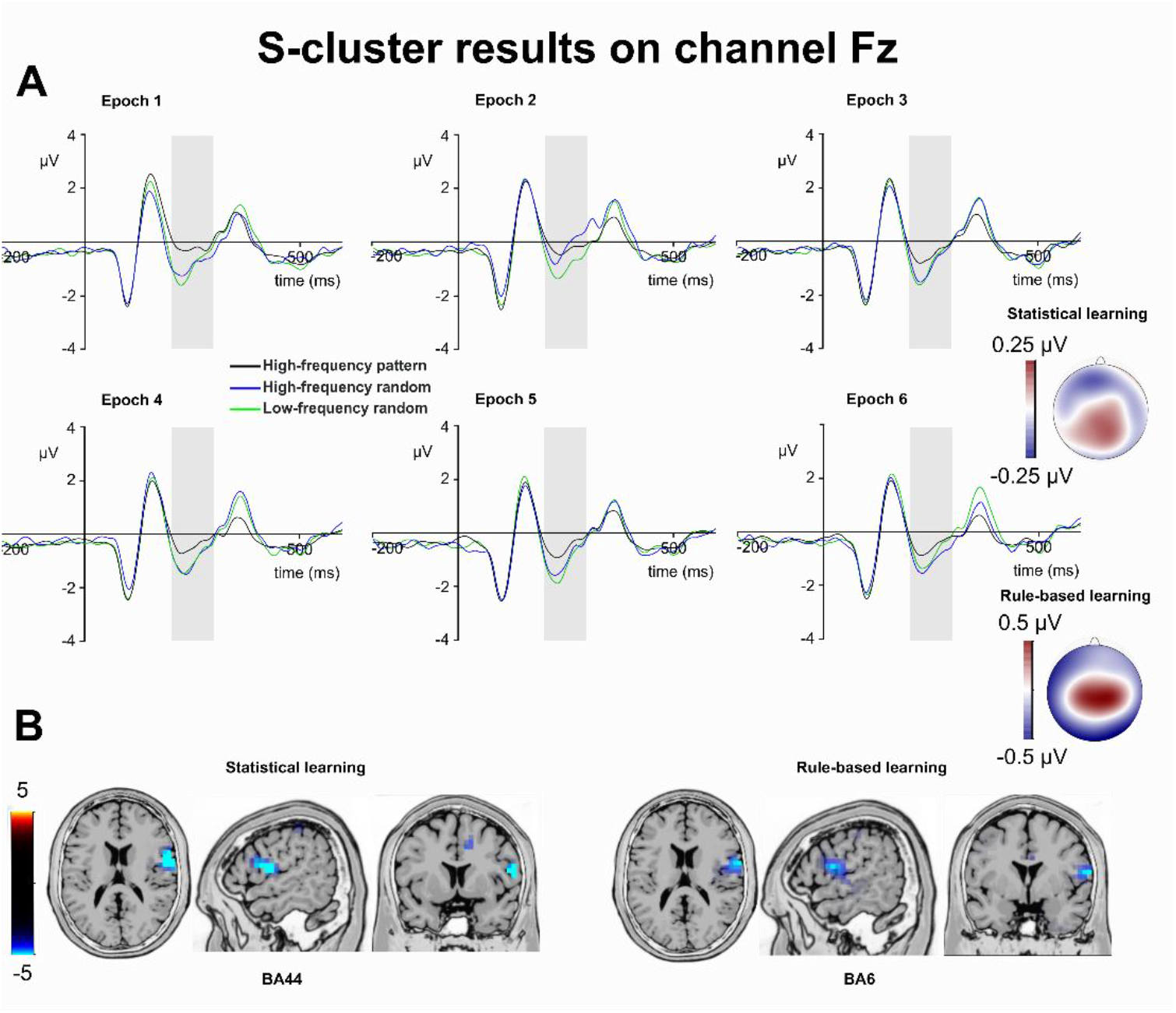
S-cluster N2. (A) S-cluster data is presented on channel Fz. Time point zero represents the stimulus presentation. The analysed time window (200-300 ms) is marked with a shaded area. The S-cluster N2 is presented across three conditions: high-frequency pattern (black), high-frequency random (blue), and low-frequency random (green). The six panels depict the six consecutive epochs of the task. The scalp topography plots show the distribution of the mean activity of the two main contrasts: statistical learning as a difference between low-frequency random and high-frequency random and rule-based learning as a difference between high-frequency pattern and high-frequency random conditions. (B) Voxels with significant differences for the statistical learning and rule-based learning effects according to the standard low resolution brain electromagnetic tomography (sLORETA) analysis are presented. The sLORETA colour bar presents critical *t* values.

**Table 1:** Summary of the decomposed N2 findings.

| Analysis and effects | S-cluster |  |  | C-cluster |  |  | R-cluster |  |  |
| --- | --- | --- | --- | --- | --- | --- | --- | --- | --- |
| | *F* | *p* | $\eta_p^2$ | *F* | *p* | $\eta_p^2$ | *F* | *p* | $\eta_p^2$ |
| *General* |  |  |  |  |  |  |  |  |  |
| Type | 10.73 | < .001 | .216 | 9.2 | .001 | .191 | 0.79 | .409 | .020 |
| Epoch | 3.97 | .002 | .092 | 1.27 | .287 | .032 | 1.58 | .169 | .039 |
| Interaction | 2.2 | .036 | .053 | 3.98 | < .001 | .092 | 0.22 | .967 | .006 |
| *Statistical learning* |  |  |  |  |  |  |  |  |  |
| Type | 1.52 | .226 | .037 | 4.54 | .039 | .104 | - | - | - |
| Epoch | 2.99 | .020 | .071 | 0.48 | .755 | .012 | - | - | - |
| Interaction | 3.14 | .016 | .074 | 1.93 | .105 | .047 | - | - | - |
| *Rule-based learning* |  |  |  |  |  |  |  |  |  |
| Type | 9.53 | .004 | .196 | 7.48 | .009 | .161 | - | - | - |
| Epoch | 5.33 | < .001 | .120 | 2.6 | .044 | .062 | - | - | - |
| Interaction | 1.98 | .101 | .048 | 3.81 | .003 | .089 | - | - | - |

The type by epoch ANOVA on the mean amplitude of the S-cluster N2 showed that the main effect of type was significant (*F*(2,78) = 10.73, ε = .809, *p* < .001, ηp^2^ = .216). Similarly, the main effect of epoch was significant (*F*(5,195) = 3.97, *p* = .002, ηp^2^ = .092). Importantly, the type by epoch interaction was also significant (*F*(10,390) = 2.20, ε = .674, *p* = .036, ηp^2^ = .053), suggesting that the mean amplitude of the S-cluster N2 changed between triplet types during the task.

In case of *statistical learning,* the type by epoch ANOVA showed that the main effect of triplet type was not significant (*F*(1,39) = 1.52, *p* = .226, ηp^2^ = .037). However, the main effect of epoch was significant (*F*(5,195) = 2.99, ε = .811, *p* = .020, ηp^2^ = .071). The S-cluster N2 was larger in the 5^th^ (−1.07 μV ± .26) than in the second epoch (−0.54 μV ± .28, *p* = .027). None of the other differences between epochs were significant *(p* > .070). However, the type by epoch interaction was significant (*F*(5,195) = 3.14, ε = .796, *p* = .016, ηp^2^ = .074). In the 2^nd^ epoch, the S-cluster N2 amplitude was larger in the low-frequency random (−0.90 μV ± .26) than in the high-frequency random (−0.19 μV ± .32, *p* < .001) condition. Triplet types did not differ from each other in other epochs *(ps* > .155). The sLORETA analysis revealed that the statistical learning effect was reflected by activation modulations in the right IFG (BA44; MNI [x,y,z]: 60, 5, 15). Thus, in the S-cluster N2, statistical learning was observed as a rapid effect occurring between the first and second epochs of the experiment, and this effect was related to right IFG activation.

In case of *rule-based learning,* the type by epoch ANOVA showed that the main effect of type was significant (*F*(1,39) = 9.53, *p* = .004, ηp^2^ = .196). The S-cluster N2 was larger in the high-frequency random (−0.83 μV ± .24) than in the high-frequency pattern (−0.41 μV ± .23) condition. Similarly, the main effect of epoch was significant (*F*(5,195) = 5.33,*p* < .001, ηp^2^ = .120). The S-cluster N2 was smaller in the 2^nd^ epoch (−0.21 μV ± .26) than in the 3^rd^ (−0.68 μV ± .23, *p* = .008), 4^th^ (−0.69 μV ± .23, *p* = .037), 5^th^ (−0.78 μV ± .24, *p* = .001), and 6^th^ epochs (−0.80 μV ± .24, *p* < .001), consecutively. None of the other epochs differed from each other *(p* > .296). The type by epoch interaction was not significant (*F*(5,195) = 1.98, ε = .796, *p* = .101, ηp^2^ = .048). According to the sLORETA analysis, the rule-based learning effect was reflected by activation modulations in the right prefrontal gyrus (BA6; MNI [x,y,z]: 65, 0, 15). Thus, rule-based learning showed time-invariant effect in the S-cluster N2 mean amplitude. Triplets ending with pattern or random elements with the same frequency were dissociated through the whole task, and this difference was related to prefrontal activity.

#### 3.2.2 Decomposed N2 (C-cluster and R-cluster)

Grand-averages of ERP waveforms in the C-cluster N2 time window split by triplet type and epoch are presented in Figure 3 and statistical results are summarized in Table 1.

**Figure 3.**
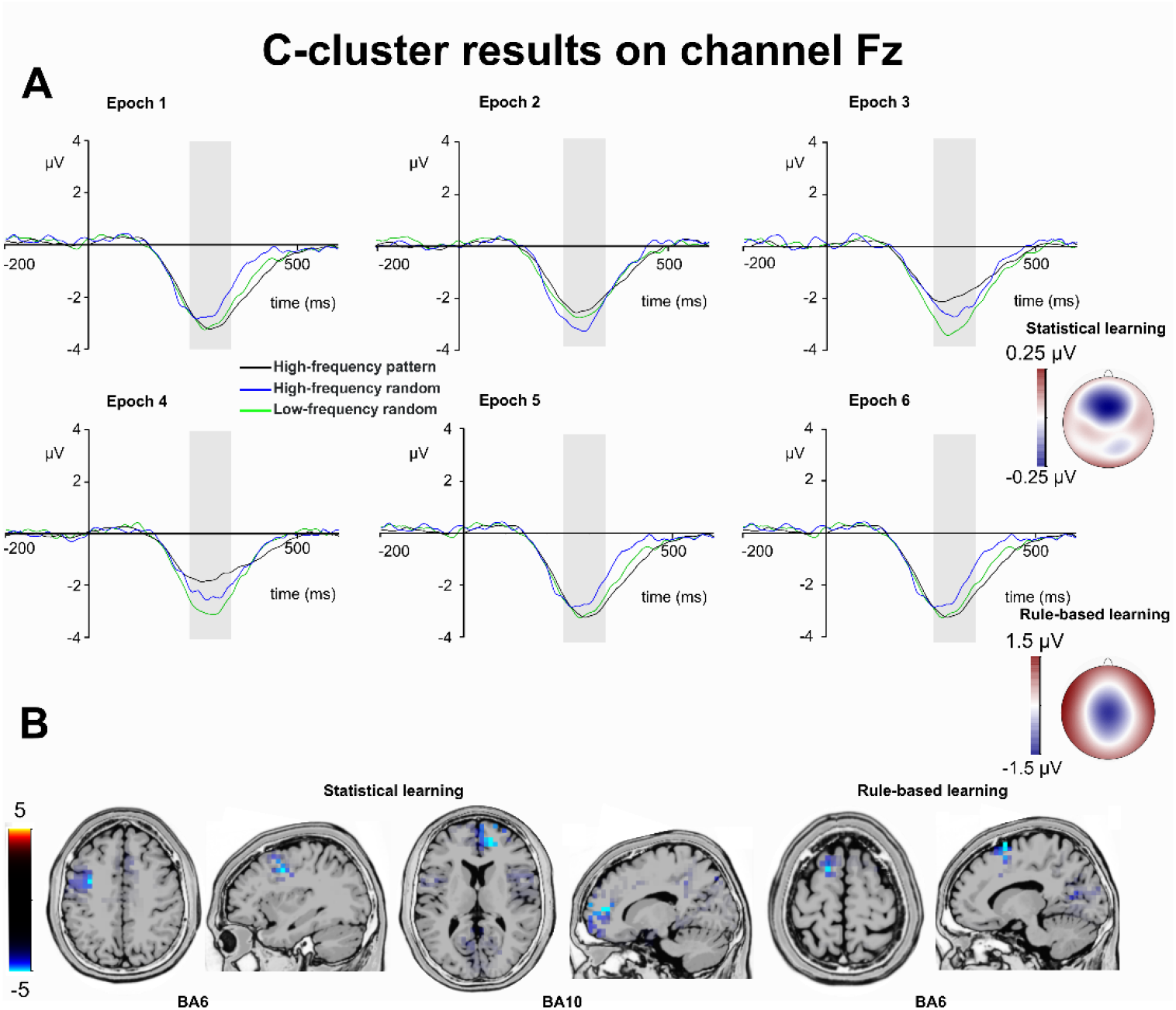
C-cluster N2. (A) C-cluster data is presented on channel Fz. Time point zero represents the stimulus presentation. The analysed time window (240-340 ms) is marked with a shaded area. The C-cluster N2 is presented across three conditions: high-frequency pattern (black), high-frequency random (blue), and low-frequency random (green). The six panels depict the six consecutive epochs of the task. The scalp topography plots show the distribution of the mean activity of the two main contrasts: statistical learning as a difference between low-frequency random and high-frequency random and rule-based learning as a difference between high-frequency pattern and high-frequency random conditions. (B) Voxels with significant differences for the statistical learning and rule-based learning effects according to the standard low resolution brain electromagnetic tomography (sLORETA) analysis are presented. The sLORETA colour bar presents critical *t* values.

The type by epoch ANOVA on the mean amplitude of the C-cluster N2 showed that the main effect of type was significant (*F*(2,78) = 9.20, ε = .715, *p* = .001, ηp^2^ = .191). However, the main effect of epoch was not significant (*F*(5,195) = 1.27, ε = .679, *p* = .287, ηp^2^ = .032). Importantly, the type by epoch interaction was significant (*F*(10,390) = 3.98, ε = .695,*p* < .001, ηp^2^ = .092), which suggests that the mean amplitude of the C-cluster N2 changed between triplet types during the task. In case of *statistical learning*, the type by epoch ANOVA showed that the main effect of triplet was significant (*F*(1,39) = 4.54, *p* = .039, ηp^2^ = .104). The C-cluster N2 was larger in the low-frequency random (−2.91 μV ± .36) than in the high-frequency random (−2.65 μV ± .33) condition. However, the main effect of epoch was not significant (*F*(5,195) = 0.48, ε = .832, *p* = .755, ηp^2^ = .012). Similarly, the triplet by epoch interaction was not significant (*F*(5,195) = 1.93, ε = .827, *p* = .105, ηp^2^ = .047). The sLORETA analysis revealed that the statistical learning effect was reflected by activation modulations in the left middle frontal gyrus (BA6; MNI [x,y,z]: −35, 0, 40) and in the right medial frontal gyrus (BA10; MNI [x,y,z]: 15, 50, 10). Thus, statistical learning effect occurred as a difference between low-frequency random and high-frequency random triplets irrespective of time on task, and this effect of the C-cluster N2 was related to prefrontal activities.

In case of *rule-based learning,* the type (high-frequency random vs high-frequency pattern) by epoch (1-6) ANOVA showed that the main effect of type was significant (*F*(1, 39) = 7.48, *p* = .009, ηp^2^ = .161). The C-cluster N2 was larger in the high-frequency random (−2.65 μV ± .33) than in the high-frequency pattern (−2.12 μV ± .33) condition. Similarly, the main effect of epoch was significant *(F(5,* 195) = 2.60, ε = .725, *p* = .044, ηp^2^ = .062). The C-cluster N2 was smaller in the 3^rd^ epoch (−2.20 μV ± .35) than in the 1^st^ (−2.78 μV ± .34, *p* = .017) and 2^nd^ (−2.66 μV ± .34, *p* = .017), and it was smaller in the 4^th^ epoch (−2.03 μV ± .40) than in the 1^st^ *(p* = .009), and 2^nd^ epochs *(p* = .015). None of the other epochs differed from each other *(p* > .082). The type by epoch interaction was also significant (*F*(5, 195) = 3.81, *p* = .003, ηp^2^ = .089). The C-cluster N2 was smaller in the high-frequency random than in the high-frequency pattern condition in the 5^th^ (−2.83 μV ± .37 vs. −2.01 μV ± .37, *p* = .019) and 6^th^ epochs (−2.78 μV ± .39 vs. −1.68 μV ± .37, *p* = .001). The difference between conditions was not significant in the other epochs *(p* > .061). The sLORETA analysis revealed that the rule-based learning effect was reflected by activation modulations in the left superior frontal gyrus (BA6; MNI [x,y,z]: −15, 10, 70). Thus, rule-based learning was detected in the C-cluster N2 mean amplitude data as a gradually increasing difference between random and pattern elements with the same frequency. This difference became the largest by the end of the learning, and it was related to activation in the superior frontal gyrus.

The type (high-frequency pattern, high-frequency random, and low-frequency random) by epoch (1-6) ANOVA on the mean amplitude of the R-cluster N2 showed that the main effects of type (*F*(2, 78) = 0.79, ε = .641, *p* = .409, ηp^2^ = .020) and epoch (*F*(5, 195) = 1.58, *p* = .169, ηp^2^ = .039) were not significant. Similarly, the type by epoch interaction was not significant either (*F*(5, 195) = 0.22, ε = .590, *p* = .967, ηp^2^ = .006). Thus, the response-related R-cluster N2 did not show any modulation related to either statistical learning or rule-based learning.

#### 3.2.3 Decomposed P3 (S-cluster)

Grand-averages of ERP waveforms in the S-cluster P3 time windows split by triplet type and epoch are presented in Figure 4 and statistical results are summarized in Table 2.

**Figure 4.**
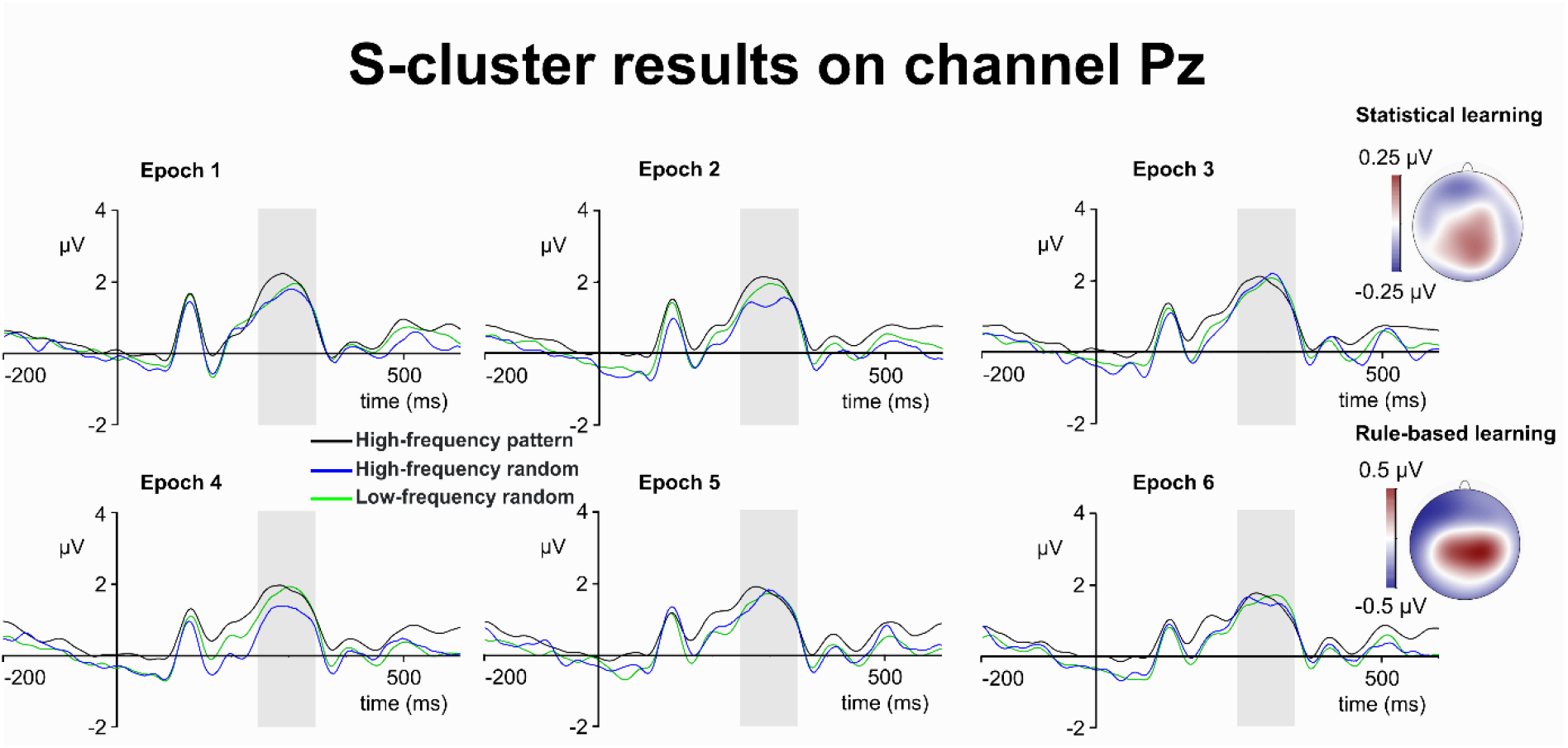
S-cluster P3. S-cluster P3. S-cluster data is presented on channel Pz. Time point zero represents the stimulus presentation. The analysed time window (250-350 ms) is marked with a shaded area. The S-cluster P3 is presented across three conditions: high-frequency pattern (black), high-frequency random (blue), and low-frequency random (green). The six panels depict the six consecutive epochs of the task. The scalp topography plots show the distribution of the mean activity of the two main contrasts: statistical learning as a difference between low-frequency random and high-frequency random and rule-based learning as a difference between high-frequency pattern and high-frequency random conditions.

**Table 2:**
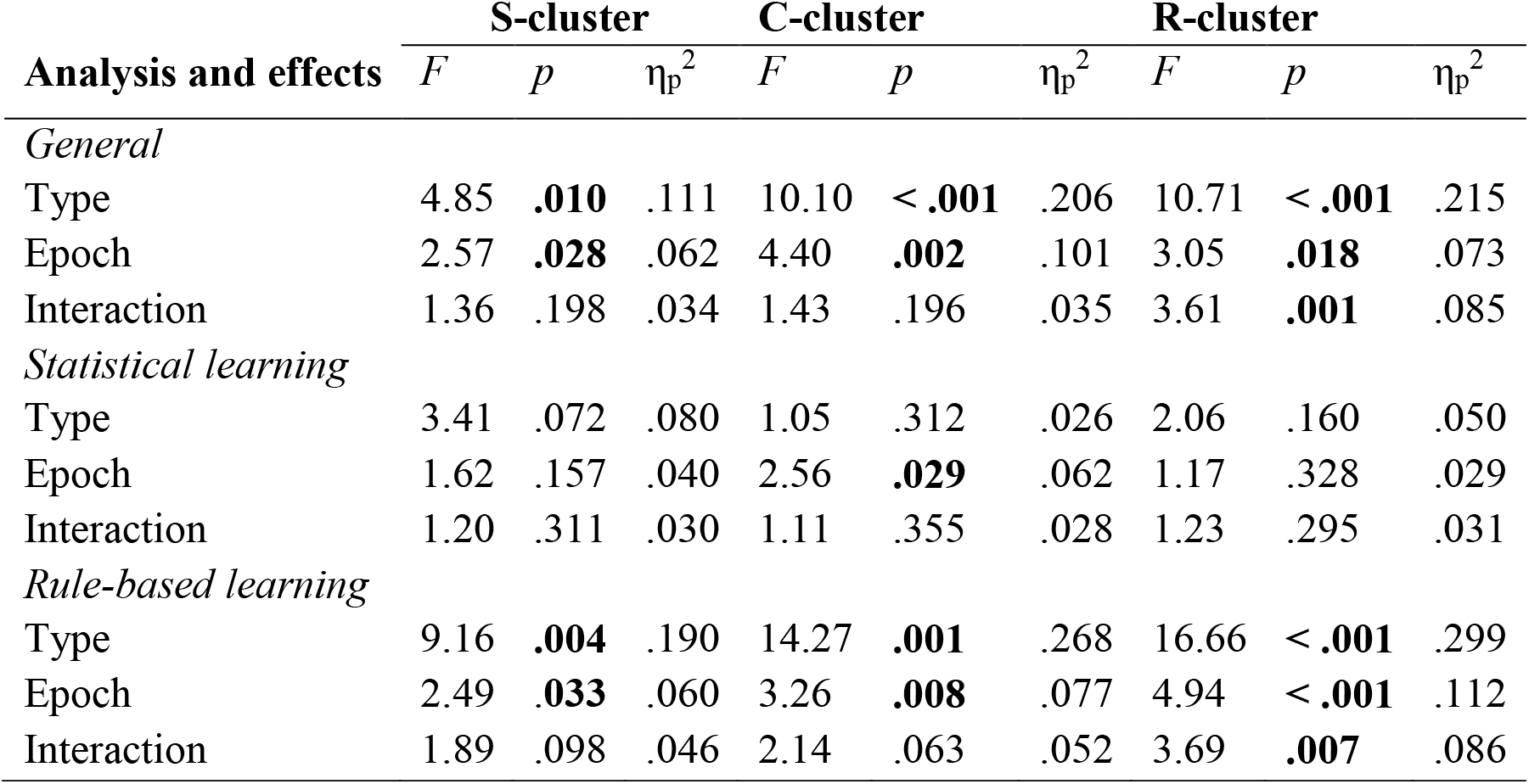
Summary of the decomposed P3 findings.

The type by epoch ANOVA on the mean amplitude of the S-cluster P3 showed that the main effect of type was significant (*F*(2,78) = 4.85, *p* = .010, ηp^2^ = .111). Similarly, the main effect of epoch was significant (*F*(5,195) = 2.57, *p* = .028, ηp^2^ = .062). However, the type by epoch interaction was not significant (*F*(10,390) = 1.36, *p* = .198, ηp^2^ = .034). In case of *statistical learning,* the type by epoch ANOVA showed that the main effects of type (*F*(1,39) = 3.41,*p* = .072, ηp^2^ = .080) and epoch (*F*(5,195) = 1.62, *p* = .157, ηp^2^ = .040) were not significant. Similarly, the type by epoch interaction was not significant (*F*(5,195) = 1.20, *p* = .311, ηp^2^ = .030). Thus, the S-cluster P3 did not show any modulation related to statistical learning. In case of *rule-based learning,* the type by epoch ANOVA showed that the main effect of type was significant (*F*(1,39) = 9.16, *p* = .004, ηp^2^ = .190). The S-cluster P3 was larger in the high-frequency pattern (1.74 μV ± .18) than in the high-frequency random (1.48 μV ± .21) condition. Similarly, the main effect of epoch was significant (*F*(5, 195) = 2.49, *p* = .033, ηp^2^ = .060). The S-cluster P3 was smaller in the 4^th^ epoch (1.47 μV ± .21) than in the 3^rd^ (1.80 μV ± .21, *p* = .023). None of the other epochs differed from each other *(p* > .402). However, the type by epoch interaction was not significant (*F*(5,195) = 1.89, *p* = .098, ηp^2^ = .046). The sLORETA analysis revealed that the rule-based learning effect was reflected by activation modulations in the left IFG (BA46; MNI [x,y,z]: −50, 10, 35) and in the left anterior cingulate cortex (BA39; MNI [x,y,z]: −5, 25, 30). Thus, the S-cluster P3 was sensitive the rule-based learning.

#### 3.2.4 Decomposed P3 (C-cluster)

Grand-averages of ERP waveforms in the C-cluster P3 time windows split by triplet type and epoch are presented in Figure 5 and statistical results are summarized in Table 2.

**Figure 5.**
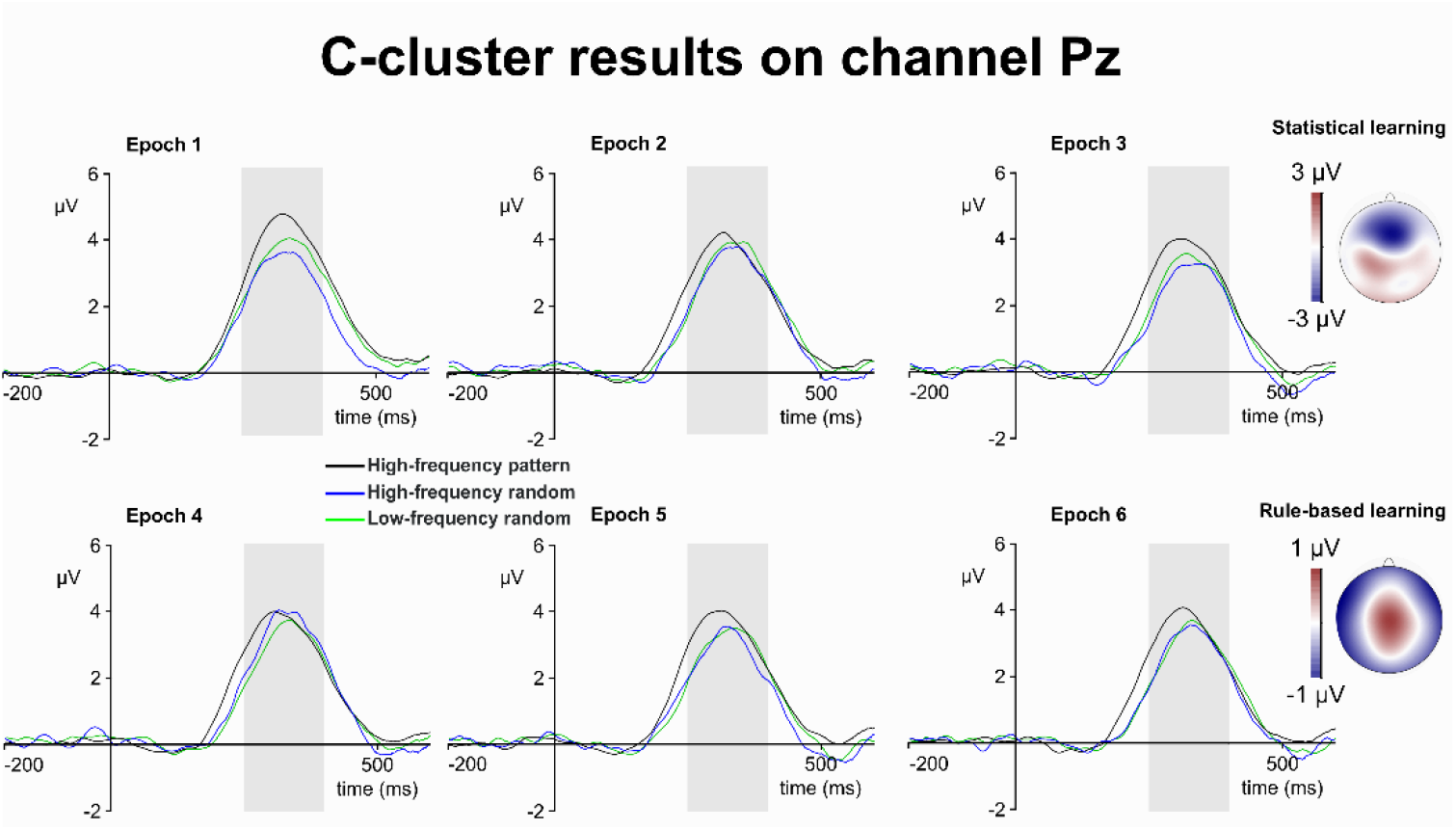
C-cluster P3. C-cluster data is presented on channel Pz. Time point zero represents the stimulus presentation. The analysed time window (250-400 ms) is marked with a shaded area. The C-cluster P3 is presented across three conditions: high-frequency pattern (black), high-frequency random (blue), and low-frequency random (green). The six panels depict the six consecutive epochs of the task. The scalp topography plots show the distribution of the mean activity of the two main contrasts: statistical learning as a difference between low-frequency random and high-frequency random and rule-based learning as a difference between high-frequency pattern and high-frequency random conditions.

The type by epoch ANOVA on the mean amplitude of the C-cluster P3 showed that the main effect of type was significant (*F*(2,78) = 10.10, ε = .862, *p* < .001, ηp^2^ = .206). Similarly, the main effect of epoch was significant (*F*(5,195) = 4.40, ε = .822, *p* = .002, ηp^2^ = .101). However, the type by epoch interaction was not significant (*F*(10,390) = 1.43, ε = .662, *p* = .196, ηp^2^ = .035). In case of *statistical learning*, the type by epoch ANOVA showed that the main effect of triplet type (*F*(1, 39) = 1.05, *p* = .312, ηp^2^ = .026) was not significant. The main effect of epoch (*F*(5,195) = 2.56, *p* = .029, ηp^2^ = .062) was significant, however, after Bonferroni-correction, none of the pair-wise differences between the epochs were significant *(ps* > .182). The type by epoch interaction was not significant either (*F*(5,195) = 1.11, *p* = .355, ηp^2^ = .028). In case of *rule-based learning,* the type (high-frequency random vs high-frequency pattern) by epoch (1-6) ANOVA showed that the main effect of type was significant (*F*(1,39) = 14.27, *p* = .001, ηp^2^ = .268). The C-cluster P3 was larger in the high-frequency pattern (3.60 μV ± .24) than in the high-frequency random (3.05 μV ± .25) condition. Similarly, the main effect of epoch was significant (*F*(5, 195) = 3.26, *p* = .008, ηp^2^ = .077). The C-cluster P3 was smaller in the 5^th^ epoch (3.15 μV ± .25) than in the 1^st^ one (3.64 μV ± .27, *p* = .028). None of the other epochs differed from each other *(ps* > .231).

The type by epoch interaction was not significant (*F*(5, 195) = 2.14, *p* = .063, ηp^2^ = .052). The sLORETA analysis revealed that the rule-based learning effect was reflected by activation modulations in the right middle frontal gyrus (BA46; MNI [x,y,z]: 50, 40, 20) and in the right middle temporal gyrus (BA39; MNI [x,y,z]: 45, −75, 10). Thus, rule-based learning modulated the C-cluster P3 mean amplitude. High-frequency pattern and random triplets were dissociated through the whole task, and this difference was related to frontotemporal activities.

#### 3.2.5 Decomposed P3 (R-cluster)

Grand-averages of ERP waveforms in the R-cluster P3 time windows split by triplet type and epoch are presented in Figure 6 and statistical results are summarized in Table 2.

**Figure 6.**
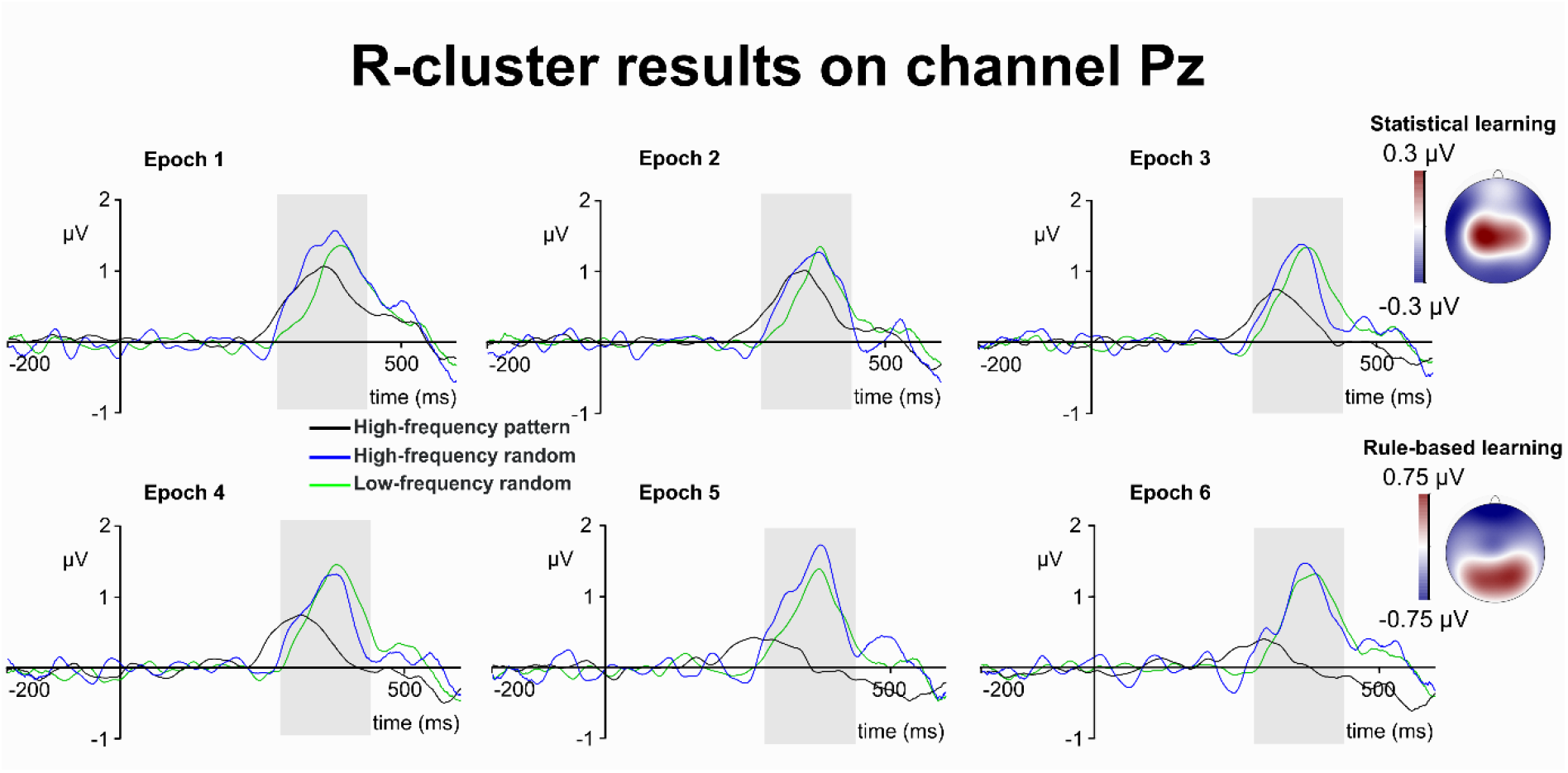
R-cluster P3. R-cluster data is presented on channel Pz. Time point zero represents the stimulus presentation. The analysed time window (280-440 ms) is marked with a shaded area. The R-cluster P3 is presented across three conditions: high-frequency pattern (black), high-frequency random (blue), and low-frequency random (green). The six panels depict the six consecutive epochs of the task. The scalp topography plots show the distribution of the mean activity of the two main contrasts: statistical learning as a difference between low-frequency random and high-frequency random and rule-based learning as a difference between high-frequency pattern and high-frequency random conditions.

The type by epoch ANOVA on the mean amplitude of the R-cluster P3 showed that the main effect of type was significant (*F*(2,78) = 10.71, ε = .853, *p* < .001, ηp^2^ = .215). Similarly, the main effect of epoch was significant (*F*(5,195) = 3.05, ε = .813, *p* = .018, ηp^2^ = .073). Importantly, the type by epoch interaction was also significant (*F*(10,390) = 3.61, ε = .649, *p* = .001, ηp^2^ = .085). In case of *statistical learning,* the type by epoch ANOVA showed that the main effects of triplet type (*F*(1,39) = 2.06, *p* = .160, ηp^2^ = .050) and epoch (*F*(5,195) = 1.17, *p* = .328, ηp^2^ = .029) were not significant. Similarly, the type by epoch interaction was not significant either (*F*(5,195) = 1.23, ε = .804, *p* = .295, ηp^2^ = .031). In case of *rule-based learning,* the type by epoch ANOVA showed that the main effect of type was significant (*F*(1,39) = 16.66, *p* < .001, ηp^2^ = .299). The R-cluster P3 was larger in the high-frequency random (0.91 μV ± .15) than in the high-frequency pattern (0.41 μV ± .12) condition. Similarly, the main effect of epoch was also significant (*F*(5, 195) = 4.94, *p* < .001, ηp^2^ = .112). The R-cluster P3 was larger in the 1^st^ epoch (0.96 μV ± .14) than in the 5^th^ (0.58 μV ± .13, *p* = .014) and 6^th^ epochs (0.49 μV ± .12, *p* = .012). None of the other epochs differed from each other *(ps* > .066). Importantly, the type by epoch interaction was also significant (*F*(5,195) = 3.69, ε = .794, *p* = .007, ηp^2^ = .086). The R-cluster P3 was larger for high-frequency random (1.13 μV ± .18) than for high-frequency random trials in the 1^st^ epoch (0.79 μV ± .14, *p* = .042). The same difference occurred in the 3^rd^ (0.78 μV ± .20 vs. 0.42 μV ± .16, *p* = .039), 4^th^ (0.81 μV ± .19 vs. 0.40 μV ± .15, *p* = .027), 5^th^ (1.05 μV ± .19 vs. 0.11 μV ± .14, *p* < .001), and 6^th^ epochs (0.87 μV ± .17 vs. 0.10 μV ± .15, *p* < .001). The two conditions did not differ from each other in the 2^nd^ epoch (*p* = .333). Thus, rule-based learning was detected in the R-cluster P3 mean amplitude. Moreover, the interaction revealed a gradually increasing difference between high-frequency random and pattern elements. This difference grew the largest by the end of the experiment, similarly to the C-cluster N2 results (see above).

## 4. Discussion

We compared temporally decomposed neurophysiological correlates of parallel learning processes, namely statistical learning and rule-based learning. The temporal decomposition successfully differentiated between S- and C-cluster activities in the time windows of the N2 component, and between S-, C-, and R-cluster activities in the time windows of the P3 component. We expected that statistical learning is reflected by the S-cluster activity, while rule-based learning is reflected by the C-cluster activity. The results partially confirmed these hypotheses; however, the difference was more gradual than categorical between the learning mechanisms. In the following sections, we discuss how statistical learning and rule-based learning effects modulated the temporally decomposed EEG signal. After reviewing the results of the RIDE clusters, we compare the two mechanisms and discuss the associated functional neuroanatomical sources.

### 4.1 Statistical learning

Statistical learning occurred incidentally, reflecting a rapid, stimulus-driven, implicit process (Kóbor et al., 2018). This was reflected by both S-cluster and C-cluster N2 activities, as random triplets were distinguishable according to their predictability (e.g. probability information) in these two clusters. Importantly, the C-cluster N2 did not show any modulation by statistical learning as the task progressed. This is in line with the results of Kóbor et al. (2018), where the N2 statistical learning effect was time-invariant, thus, it did not show a gradual accumulation of statistical knowledge. However, the statistical learning effect of the S-cluster N2 was time-variant, as the difference between triplet types increased between the first and second epochs. Notably, this rapidly occurring effect decreased by the next epoch, which then showed similar activity to the beginning of the task. Thus, changes in the S-cluster N2 were rapid and short lasting, while the C-cluster N2 statistical learning effect was constant during the task. This is in line with the behavioural results that showed a rapid acquisition of statistical information early in the task, which then remained stable during the experiment (Kóbor et al., 2018). Both the C-cluster N2 and the S-cluster N2 mean amplitudes were larger for low-frequency random than for high-frequency random triplets. Thus, less predictable stimuli triggered larger N2 responses both as a function of mismatch or novelty detection (S-cluster) and as a signal of response conflict (C-cluster). Detecting statistical learning effects both in S-cluster N2 and C-cluster N2 promotes the notion of Koelsch et al. (2016) that processing transitional probabilities goes beyond sensory capacities. However, it is likely that a rare sequence of events signals a potential need for a higher cognitive load (Friston, 2010; Friston and Kiebel, 2009; Koelsch et al., 2016). This explanation also fits the current study, in which low-frequency random triplets were associated with slower responses, thus the stimuli that was the hardest to predict increased the processing load (Kóbor et al., 2018). This increased load was detected not only in the undecomposed N2 (Kóbor et al., 2018) but also in the C-cluster N2. It has been suggested, that a neurophysiological marker for statistical learning inevitably has a dual role: it can signal both increased processing load and a prediction error signal (Koelsch et al., 2016). Importantly, temporal signal decomposition could provide a meaningful distinction between these roles, as the S-cluster reflects the stimulus-driven aspects of novelty or mismatch detection, and the C-cluster reflects the increased cognitive load and response conflict. Thus, statistical learning operates at a perceptual level, but not exclusively. Central aspects of statistical learning likely provide signals for the higher-order functions for adaptive behaviour (Conway, 2020). Unlike previous approaches with the undecomposed EEG (Kóbor et al., 2018; Koelsch et al., 2016), the temporally decomposed clusters could differentiate between these two functions of statistical learning. Even though these simultaneous processes occur in similar and overlapping time windows, the differences between them in latency variability reveal that one is directly triggered by the visual presentation of the stimulus while the other one represents a more variable mental chronometry. Thus, the multifaceted role of learning of probabilistic information was confirmed at the neurophysiological level.

### 4.2 Rule-based learning

While statistical learning effects were specific to the S-cluster and C-cluster N2s, rule-based learning was detected in a wide range of temporally decomposed components, such as the S-cluster and C-cluster N2s, S-cluster, C-cluster, and R-cluster P3s. This pervasive effect likely indicates that the global integration of the acquired sequential information involves sensory, translational, and motor aspects concurrently. That is, rule-based learning provides a general access to summary statistics. This includes the distribution of local statistics which originates from statistical learning, and a general knowledge of the global entropy or uncertainty of the information stream (Conway, 2020; Daikoku, 2018). In the following, we discuss how temporal signal decomposition helped to differentiate between these set of functions.

Curiously, the S-cluster N2 showed a time-insensitive effect of rule-based learning. The component was larger for high-frequency random than for high-frequency pattern triplets. Thus, despite the same level of stimulus probability, sequential position triggered a stimulus-driven mismatch response. However, this effect has to be taken with caution, since in this version of the paradigm, random and pattern elements were visually distinguishable from each other, marked by red and black colours, respectively. Furthermore, rule-based learning was detected in the C-cluster N2, as well. High-frequency random stimuli which is harder to predict elicited a larger component compared to the easy to predict high-frequency pattern condition. Unlike statistical learning, the rule-based learning effect on the C-cluster N2 increased with practice, showing a gradual accumulation of sequence knowledge. This is in line with the undecomposed N2 results (Kóbor et al., 2018) and earlier reports of ERP correlates of sequence learning (Eimer et al., 1996; Ferdinand et al., 2007; Rüsseler et al., 2003). The time-sensitive nature of rule-based learning also reflects the behavioural results that showed continuous learning as the task progressed (Kóbor et al., 2018). Thus, the C-cluster N2 showed evidence for the changing level of response conflict between predictable (learnt) and less predictable random triplets.

Rule-based learning effects were detected in the decomposed P3 components, as well. Effect of attention on detecting irregularities in the information stream is traditionally linked to the P3 (Chennu and Bekinschtein, 2012; Kóbor et al., 2019, 2018). Regularities, such as sequential patterns evoke attention (Zhao et al., 2013). Importantly, in a sequence in which pattern and random elements alternate each other and visually distinguishable, attention can be both endogenous and exogenous (Zhao et al., 2013). Exogenous attention evoked by the different colours of sequential and random elements is likely captured by the rule-based learning effect in the S-cluster P3. Moreover, endogenous attention evoked by internal goal might be reflected by the C-cluster P3 effect. Prioritizing a regular stimulus event over irregular ones is not a strictly stimulus-driven process since it relies on the internal representation of the regularities (Zhao et al., 2013). Interestingly, both the S-cluster P3 and the C-cluster P3 effects were constant: high-frequency pattern elements (targets according to the goal of the task) evoked larger P3s than high-frequency random elements through the task. However, this time-invariant nature of the rule-based learning effects contradict the gradual learning curve seen in the behavioural data (Kóbor et al., 2018). This contradiction was resolved by the R-cluster results. Importantly, the R-cluster P3 provided a type by epoch interaction, which showed a growing difference between sequential and random triplets with the same stimulus frequencies. That is, neurophysiological dynamics similar to the behavioural learning curve was observed in the C-cluster N2 and R-cluster P3. Thus, these two decomposed components reflected the adaptation to the sequential regularities through response conflict and response preparation mechanisms. In sum, the pervasive effect of rule learning on the temporally decomposed clusters confirm the complex nature of this learning mechanism (Conway, 2020). The learning of higher-order sequential information includes selective attention to different stimulus categories (S-cluster P3), goal-directed effort allocation (C-cluster P3), and gradually more effective response management (R-cluster P3). These various functions highlight the main characteristics of rule-based learning. Namely, it is an attention-dependent system which serves as a gating or control mechanism over the learning of sequential information (Conway, 2020).

### 4.3 Comparison between statistical learning and rule-based learning

The main goal of the current study was to differentiate between parallel learning mechanisms in the neurophysiological signal. To characterise the differences between the types of learning at a single trial level, we employed temporal signal decomposition. This was successful as statistical learning was detected specifically in the mean amplitudes of the S-cluster N2 and C-cluster N2, while rule-based learning effect occurred in the mean amplitudes ranging from the S-cluster N2 to the R-cluster P3. Thus, statistical learning was identified as a learning mechanism related to mismatch-detection and early response conflict. In contrast, rule-based learning as a higher-order process was reflected by every aspect of the decomposed N2 and P3 signals except of the R-cluster N2. Thus, it was confirmed that rule-based learning builds upon the acquired statistical regularities, and contributes to the control of the learning (Conway, 2020). However, a specific statistical learning effect and a more general rule-based learning also raises the possibility that the two learning mechanisms are the same process albeit with different levels of complexity. This alternative explanation was ruled out by the source localization analyses. An important validation whether the decomposed components are related to distinct processes is differentiating between their neural sources. Specifically, we have expected that statistical learning effect of the the S-cluster N2 is associated with right IFG activity, while rule-based learning as reflected by the C-cluster N2 is associated with a more widespread dorsolateral activity. This was partially confirmed by the source localization, which revealed that the statistical learning effect on the S-cluster N2 was related to activation modulations in the right IFG. In contrast, rule-based learning effect was reflected by the right prefrontal gyrus’ activation. The C-cluster N2 also showed differences in the neural sources of the two forms of learning: statistical learning effect was reflected by changes of activation in the left middle frontal gyrus and in the right medial frontal gyrus, while rule-based learning was reflected by activation modulations in the left superior frontal gyrus. Thus, the right IFG was mainly implicated in statistical learning, providing further evidence about this region’s importance in acquiring probabilistic information (Barascud et al., 2016; Conway, 2020; Maheu et al., 2020; Southwell and Chait, 2018). Furthermore, this association was specific for the S-cluster, while the C-cluster N2 showed more widespread prefrontal activations both for statistical and rule learning. This pattern is in line with earlier research showing that frontal sources are associated with the learning of hierarchical sequential information (Southwell and Chait, 2018). In the current study, the alternation between sequential and random items corresponds to such hierarchical structure (Nemeth et al., 2013a). Taken together, the different coding levels in the N2 time window during the task are also related to different functional neuroanatomical structures. Moreover, similar to the N2 results, the S- and C-cluster sources also differed in the P3 time windows. Please note that the P3 was only sensitive to the rule-based learning effect. This effect in the S-cluster showed activation modulations in the left IFG and in the left anterior cingulate cortex. In contrast, the C-cluster’s activity indicated the right middle frontal gyrus and the right middle temporal gyrus. In sum, different sources related to statistical learning and rule-based learning effects suggest that sequence learning does not rely on a single mechanism of predicting the upcoming stimuli. Rather, this distinction in neural sources implicates that different levels of predictions are organized in different stages of the processing hierarchy (Southwell and Chait, 2018). Thus, statistical learning and rule-based learning can be distinguished at the neurophysiological level. Additionally, the difference in neural sources does not only imply the existence of parallel processing mechanisms whilst learning a visual sequence. The specific contribution of the right IFG to the statistical learning effect of the S-cluster N2 also suggests that this region is not specific to inhibitory functions. Rather, stimulus-driven mismatch detection and perception of uncertainty of the environment may characterize the right IFG (Barascud et al., 2016; Erika-Florence et al., 2014; Southwell and Chait, 2018).

### 4.4 Conclusion

In summary, we successfully identified functionally distinguishable clusters of neurophysiological activity in the N2 and P3 time range in sequence learning. The current analyses deepen our understanding how humans are capable of learning multiple types of information from the same stimulus stream in a parallel fashion. We have demonstrated that concomitant, but distinct aspects of information coded in the N2 time window play a role in these mechanisms: mismatch detection and response control underlie statistical learning and rule-based learning, respectively, albeit with different levels of time-sensitivity. Moreover, the two learning effects in the different temporally decomposed clusters of neural activity also differed from each other in neural sources. Importantly, the right inferior frontal cortex (BA44) was specifically implicated in statistical learning, confirming its role in the acquisition of transitional probabilities. In contrast, rule-based learning was associated with the prefrontal gyrus (BA6). The results show how parallel learning mechanisms operate at the neurophysiological level and are orchestrated by distinct prefrontal cortical areas. Understanding the neural mechanisms behind primary information processing functions, such as statistical and rule-based sequence learning is crucial to gain a close-up picture of how simultaneous learning develop both in typical and atypical cognition.

## Acknowledgements

This research was supported by the Deutsche Forschungsgemeinschaft TA 1616/2-1 (to A. T.); National Brain Research Program (project 2017-1.2.1-NKP-2017-00002); Hungarian Scientific Research Fund (NKFIH-OTKA FK 124412, PI: A. K., NKFIH-OTKA K 128016, PI: D. N., NKFIH-OTKA PD 124148, PI: K. J.); János Bolyai Research Scholarship of the Hungarian Academy of Sciences (to K. J. and A. K.); IDEXLYON Fellowship of the University of Lyon as part of the Programme Investissements d’Avenir (ANR-16-IDEX-0005) (to D. N); Deutsche Forschungsgemeinschaft FOR 2698 (to C. B.).

